# Forecasting tumor growth kinetics and hypoxia levels in mice using mathematical modelling

**DOI:** 10.64898/2026.01.22.700997

**Authors:** Nicolas Azzopardi, David Ternant, Julien Sobilo, Sharuja Natkunarajah, Stéphanie Lerondel, Alain Lepape, Sébastien Roger, Stéphanie Chadet

## Abstract

Quantitative description of tumor growth is challenged by vascular heterogeneity and hypoxia. In this study, mammary tumor growth was investigated in mouse model using caliper and ultrasound imaging measurements, bioluminescence imaging (BLI), and pimonidazole imaging. Tumor volume increased monotonically when assessed by caliper and ultrasound imaging, whereas BLI exhibited oscillating dynamics despite continued tumor growth, consistent with hypoxia-related signal attenuation.

A mathematical compartmental model was developed primarily to describe tumor growth dynamics, incorporating latent vascular capacity as a key regulatory variable. The model accounts for reciprocal interactions between tumor expansion and vascular limitation. BLI was integrated as an auxiliary observable to reveal hypoxia-driven modulation of signal production rather than as a direct surrogate of tumor size. Model parameters were estimated using nonlinear mixed-effects modelling with population approach, allowing quantification of population-level behavior and inter-individual variability.

The model adequately described tumor growth while explaining BLI dynamics through vascular and hypoxic effects. This framework provides the first semi-mechanistic description of tumor growth and supports the use of BLI as an indirect marker of hypoxia and, consequently, of tumor growth. This model may provide a useful framework for quantifying the effects of vascular-modulating therapeutics and genetic polymorphisms involved in tumor progression.

## Introduction

Cancer remains one of the leading causes of morbidity and mortality worldwide and is characterized by the uncontrolled growth and spread of malignant cells. In 2025, an estimated 20 million new cancer cases were diagnosed worldwide, resulting in more than 10.3 million deaths. The complexity and heterogeneity of cancer – both between patients and within individual tumors - necessitate sustained research efforts aimed at understanding its fundamental biology, identifying novel therapeutic targets, and developing more effective treatment strategies^1^.

Cancer research has evolved substantially and relies heavily on relevant models to investigate disease progression and evaluate therapeutic interventions. These include *in vitro, ex vivo*, and *in vivo* systems, which enable the study of tumor biology under controlled conditions^2,3^. Among i*n vivo* approaches, murine models engrafted with human or mouse tumor cells (xenograft and syngeneic models, respectively) offer powerful tools to investigate tumor progression, metastatic spread, and responses to therapy^4^.

A defining feature of solid tumors is the presence of hypoxic microenvironments, in which oxygen levels fall below 5% as rapid cellular proliferation outpaces vascular supply^5^. Hypoxia constitutes a powerful selective pressure that promotes aggressive tumor phenotypes, including metabolic reprogramming, epithelial–mesenchymal transition, resistance to chemotherapy and radiotherapy, increased invasiveness, and genomic instability^6,7^.

The emergence of non-invasive imaging techniques in these models has further advanced cancer research by enabling longitudinal, real-time monitoring of tumor dynamics without the need to sacrifice experimental animals^8^. Among these modalities, bioluminescence imaging (BLI) - which relies on light emission from luciferase-expressing cells - has emerged as a powerful tool for tracking tumor growth and treatment response *in vivo*^9^. BLI offers high sensitivity, cost-effectiveness, and the ability to monitor tumor burden, proliferation, and metastasis over time in live animal models^10^.

However, the luciferase-catalyzed reaction underlying BLI is inherently oxygen-dependent; consequently, reduced oxygen availability in hypoxic tumor regions leads to attenuation of the bioluminescence signal. This quenching effect can result in a marked underestimation of tumor burden and viable cell number in severely hypoxic areas^11^. As a result, decreases in BLI signal intensity may reflect not only therapeutic efficacy but also increased tumor hypoxia, thereby confounding data interpretation and compromising quantitative accuracy^12^.

On the other hand, this oxygen dependence of BLI may also provide an indirect and informative readout of tumor hypoxia. But this relationship between tumor growth and hypoxia requires a mechanistic mathematical model to enable quantitative assessment of both processes.

In this work, we propose a new mathematical model that describes the hypoxia-induced modulation of tumor growth revealed by bioluminescence kinetics of transfected tumor cells in mice. Such a mathematical model, developed in the absence of therapeutic intervention, could serve as a foundational framework for quantifying the effects of therapeutic interventions.

## Methods

### Cells and Cell Culture

Murine mammary cancer cell line 4T1 from the Balb/cJ strain was purchased from LGC Standards (France), and a stable 4T1-luc cell line expressing the luciferase gene (thereafter called “4T1 cells”) was obtained by transduction of cells with lentiviral vectors containing the luciferase gene and blasticidin resistance gene for selection (GIGA Viral Vectors, Belgium).

### In Vivo Mammary Cancer Model and Experiments

All experiments have been approved by the Comité d’éthique du Centre-Val de Loire and have been performed in accordance with the European Ethics rules (Ref 005377.01 Apafis #12960). All animals were bred and housed at the PHENOMIN-TAAM-CNRS UAR44 (CNRS Campus, Orléans, France), in controlled conditions with a 12-hour light/dark cycle at 22°C, and free to food and water ad libitum. We used a syngeneic and orthotopic mouse mammary cancer model in female BALB/cJ immunocompetent mice. To do so, 4T1-luciferase-expressing mouse mammary cancer cells were injected into the fifth mammary fat pad of 6 weeks-old mice, as previously described^13^.

### Tumor size measurements

#### Caliper

Primary tumor size was measured with a calliper, from day 7 to day 45, every 3 to 4 days, for a total of 10 measurements. The tumor volume (mm^3^) was extrapolated as (L × W^2^)/2, where L is the length and W the width.

#### Ultrasound imaging

Tumor volume was estimated by ultrasonography, from day 7 to day 45, every 3 and 4 days, alternatively, for a total of 10 measurements.

#### Bioluminescence

Bioluminescence imaging (BLI) was performed on an IVIS-Lumina II imaging system (Perkin Elmer) generating a pseudo-colored image representing light intensity and superimposed over a greyscale reference image. Each mouse was intraperitoneally injected with luciferin potassium salt at a dose of 100 mg/kg (purchased from Promega). Mice anesthetized by 1.5% isoflurane were placed in a supine position on a thermostatically controlled heating pad (37°C) during imaging. Acquisition binning and duration were set depending on tumor activity. Signal intensity was quantified as the total flux (photons/seconds) within region of interest (ROI) drawn manually around the tumor area using Living Image 4.4 software (Perkin Elmer). BLI was performed, from day 7 to day 45, every 3 and 4 days, alternatively, for a total of 10 imaging.

### Analysis of tumor hypoxia

Pimonidazole immunostaining was performed with the Hypoxyprobe kit (Hypoxyprobe, Burlington, USA), following the supplier’s instructions. Briefly, mice received an intravenous injection of 60 mg/kg of pimonidazole solution. One hour after injection, mice were sacrificed, and tumors were resected. Tumors were then fixed in 10% formaldehyde, embedded in paraffin and sectioned. Sections were incubated overnight with anti-pimonidazole rabbit antibody, washed three times in PBS, and incubated for 1 hour at room temperature with TRITC-conjugated goat anti-rabbit antibody (Abcam, Cambridge, USA). Pimonidazole-stained tissue sections were digitalized at a high resolution for further manipulation using a Hamamatsu NanoZoomer slide scanner (Hamamatsu Photonics, Japan). Hypoxia analysis was performed by processing pimonidazole-labeled images into 8-bit format followed by binary thresholding in

ImageJ (NIH, USA). Total tissue areas, defined as regions of interest (ROIs), were identified using DAPI staining. The hypoxic extent was then expressed as the area fraction, calculated as the ratio of the pimonidazole-positive surface area to the total ROI area.

### Mathematical Model

A mathematical model was used to describe the influence of hypoxia on mammary tumor growth over time (Figure 1). The integrative mathematical model structure is presented in four parts for clarity:

**Figure 1.**
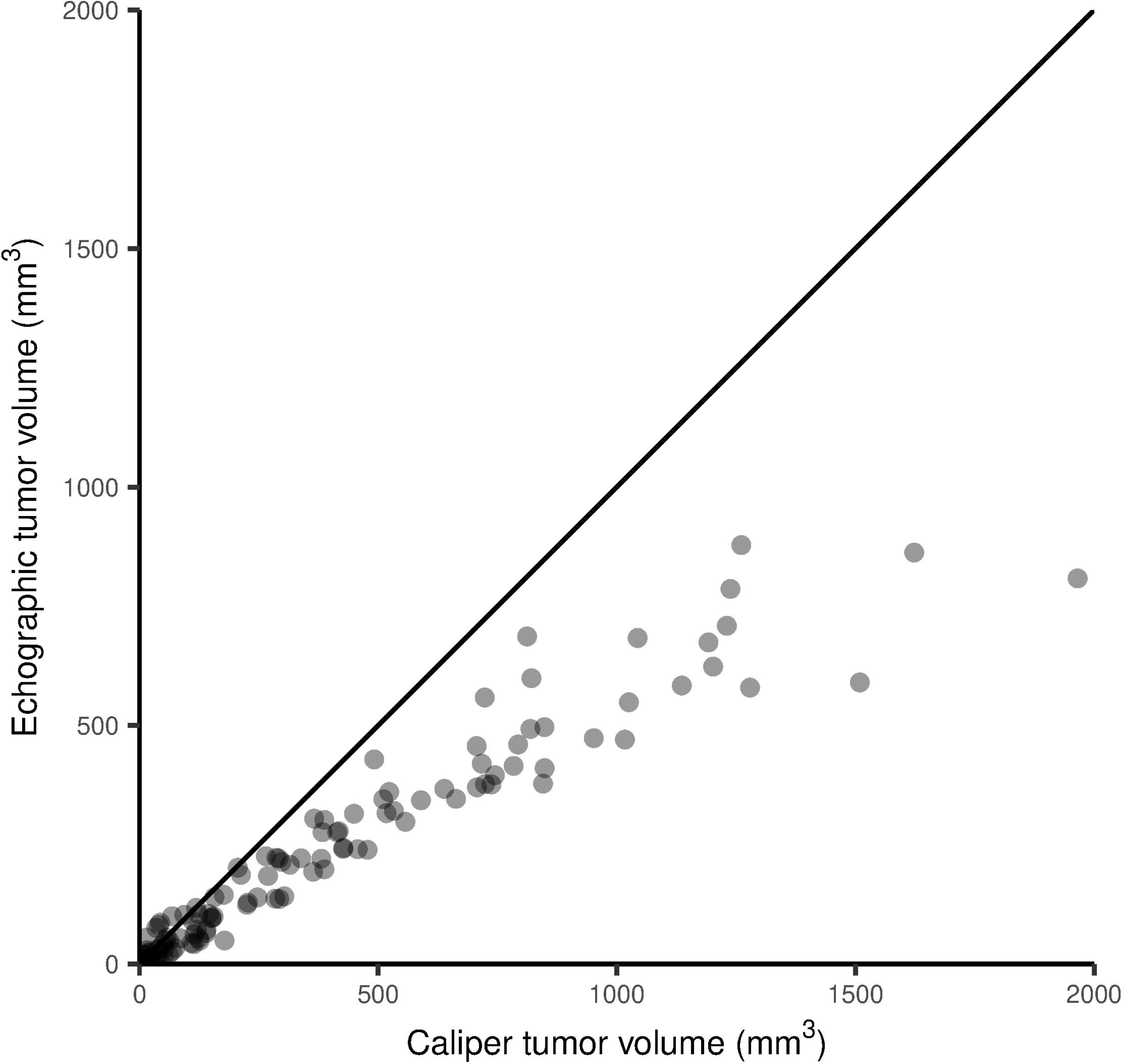
Caliper against ultrasound tumor measurement. Tumor volumes measured by caliper against ultrasound tumor volumes for matched time points. Each point corresponds to a paired measurement. Overall, caliper-based measurements tended to yield higher tumor volume estimates than ultrasound-based measurements, particularly for larger tumors.

#### Tumor growth

The injected cancer cells are represented by the initial tumor size parameter 10^*TSini*^ (mm^3^) within compartment TS. The tumor size in TS increases at growth rate *G* (mm^3^.day^-1^).

Which depends on both vascular capacity and the current tumor size (see below).

#### Vascular capacity

The estimation of the vascular capacity K is latent, as it is not directly observed. It is inferred from the dynamics of tumor size and bioluminescence signal. K is generated through a transit of three compartments characterized by a mean transit time *MKT* (day), resulting in a transfer rate between compartments of k_kt_ = (3+1)/*MKT*. The “production” and “elimination” of vascular capacity are both assumed to be k_kt_. The initial steady state vascular capacity *K*_*0*_ is modulated by the influences of vascular capacity and TS (see below).

#### Cell bioluminescence

The bioluminescence of tumor cells (BLI) is defined as a function of TS according to BLI = sL.TS.10^*gain*/10^, where sL represents the combined influence of K and TS (see below) and *gain* (dB) acts as a scaling factor that links tumor size to the intensity of bioluminescence signal.

#### Reciprocal influences

The three aforementioned influences are described by sigmoid function of 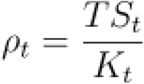, where TS_t_ and K_t_ are the Tumor size and vascular capacity, respectively, at time t.

Influence on tumor growth is 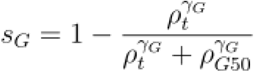, where *ρG*50 is the value of *γ*_*t*_ leading to half of growth rate G and *γ*_*G*_ is the sigmoidicity parameter.

Influence on vascular capacity production is 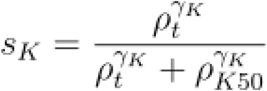, where *ρK*50 is the value of *ρK*50 leading to half of growth rate G and *γ*_*K*_ is the sigmoidicity parameter.

Influence on bioluminescence is 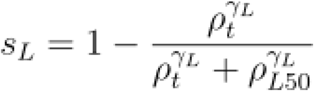, where *ρG*50 is the value of *ρ*_*t*_ leading to half of growth rate G and *γ*_*L*_ is the sigmoidicity parameter.

Model parameters were estimated using nonlinear mixed-effects modeling with population approach, which allows both population-level means and inter-individual variability characterization. This analysis was performed using Monolix 2024R1 (Simulations Plus). The complete model script is provided in supplementary materials.

## Results

### Comparison between caliper and ultrasound imaging to assess tumor size

Recognizing the potential for volumetric error when assessing irregularly shaped masses, we performed a direct comparison of standard measurement methods. Caliper-based measurements gave higher tumor size estimates than ultrasound imaging, particularly for larger tumors (Figure 1). The ultrasound measurement technique was selected, as it provides more accurate size quantification for atypically shaped tumors. Tumor size over time (TS), measured by ultrasound, shows a rapid initial increase followed by a slower growth phase.

### Variability in BLI measurements

Although bioluminescence imaging (BLI) is a powerful tool for tumor detection and longitudinal monitoring, it is inherently affected by substantial variability, which represents a major limitation. Tumor BLI exhibits an early pronounced peak, followed by a decrease that coincides with the tumor slower growth phase start. Subsequently, the BLI signal displays oscillatory fluctuations. The description and quantification of the physiological relationship between TS and BLI needs a semi-mechanistical mathematical model integration both TS and BLI observations.

### Fitting the data with the mathematical model

The mathematical model (Figure 2), which integrates the vascular capacity (K), described adequately both TS and BLI dynamics (Figure 3). The model also displayed good accuracy as shown by good concordance between observed and individual-predicted values in Figure 4A and 4B and by the normality of the Normalized Prediction Distribution Errors (NPDE) (Figures 4C and 4D).

**Figure 2.**
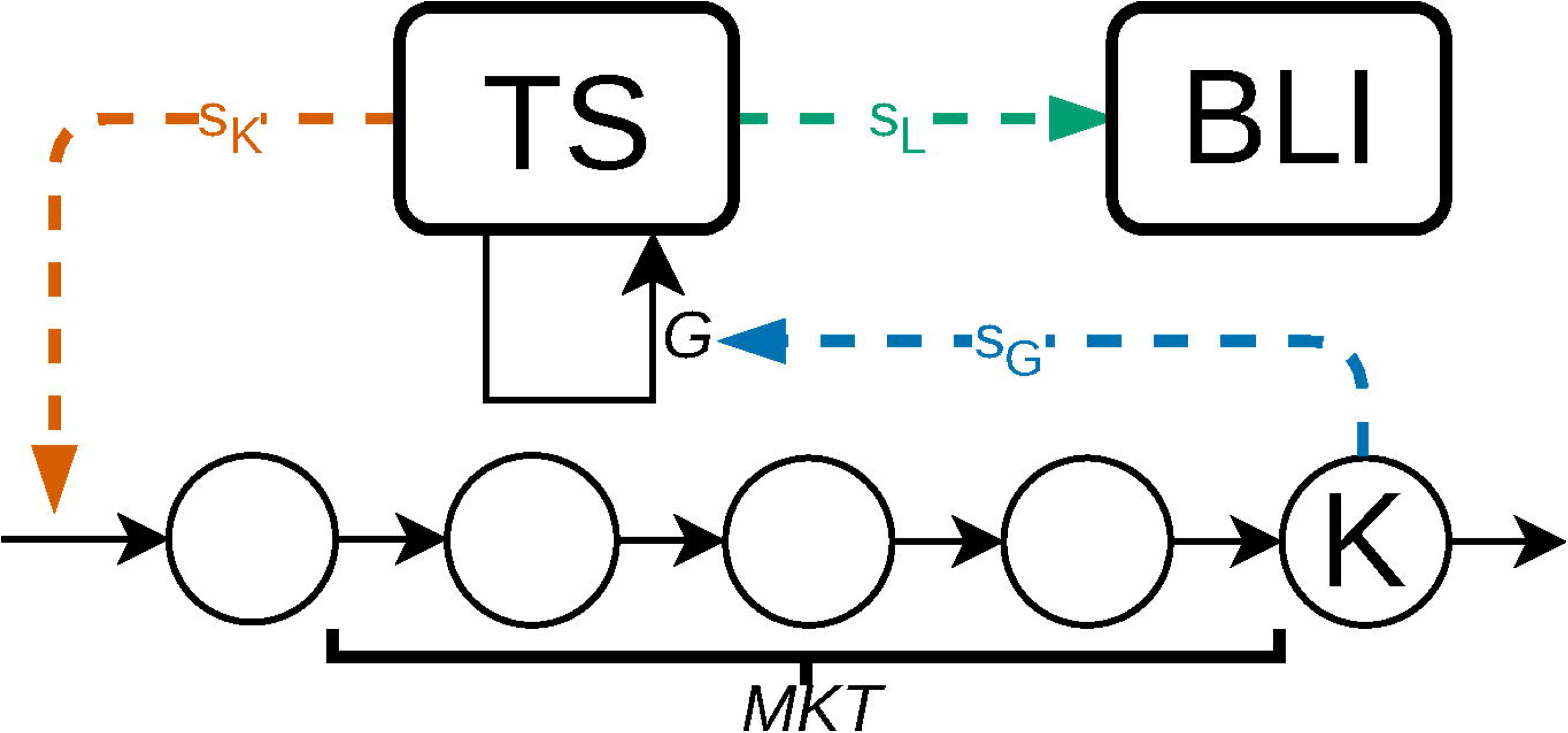
Schematic representation of the mathematical model. TS: Tumor size of growth rate *G*, L: Tumor bioluminescence; K vascular capacity of mean transit production *MKT*. Colored arrows represent the influence of TS/K ratio ρ on growth rate sG (blue), vascular capacity sK (red) and bioluminescence signal sL (green).

**Figure 3.**
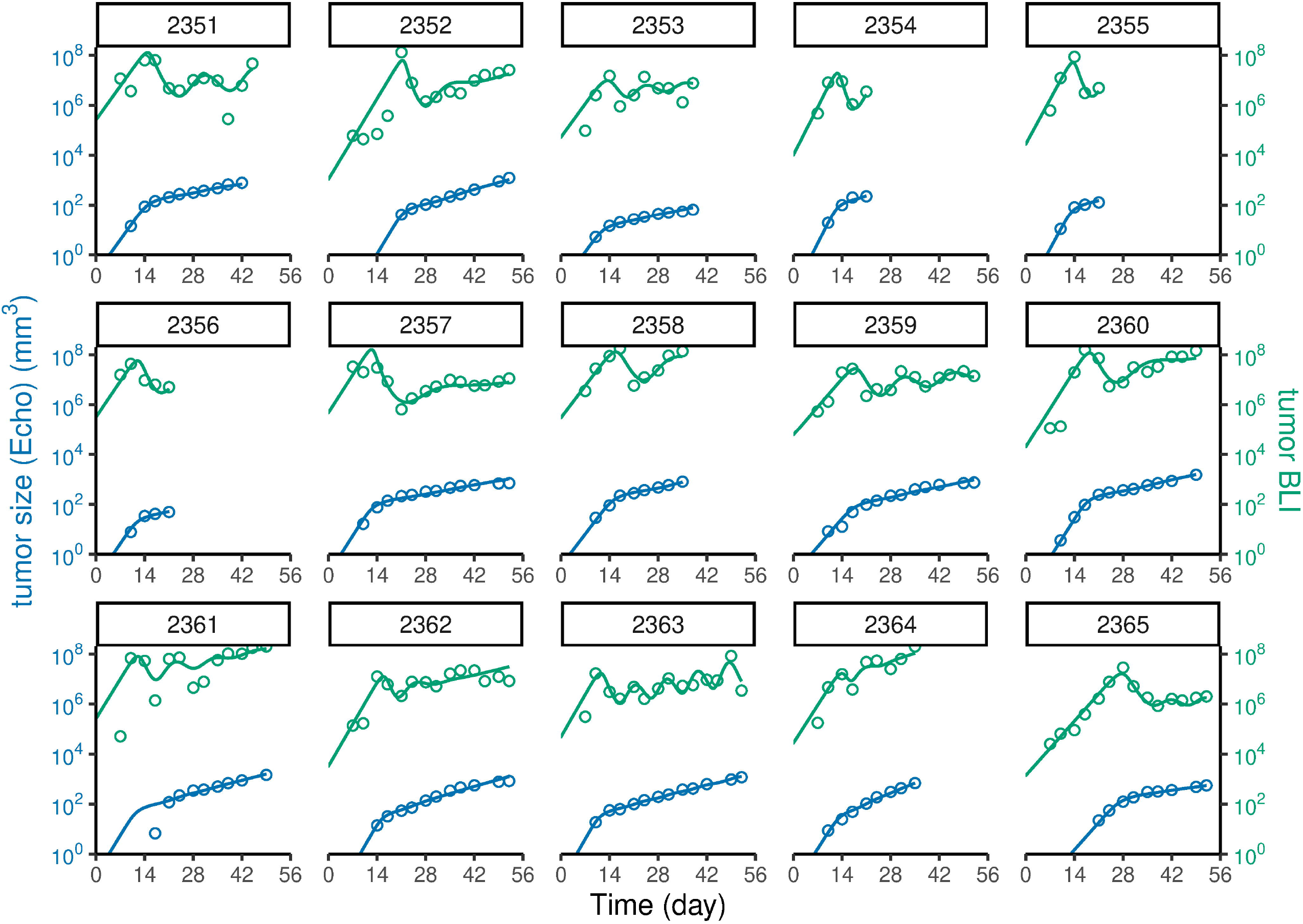
Individual profiles. Individual tumor size (blue) and bioluminescence (green) profiles of the 15 mice: - The observed values are represented by dots - The mathematical model describes correctly both tumor size and bioluminescence signal dynamics (line). After an initial rapid growth of tumor size and bioluminescence signal, the tumor growth rate starts to decrease concomitantly to bioluminescence.

**Figure 4.**
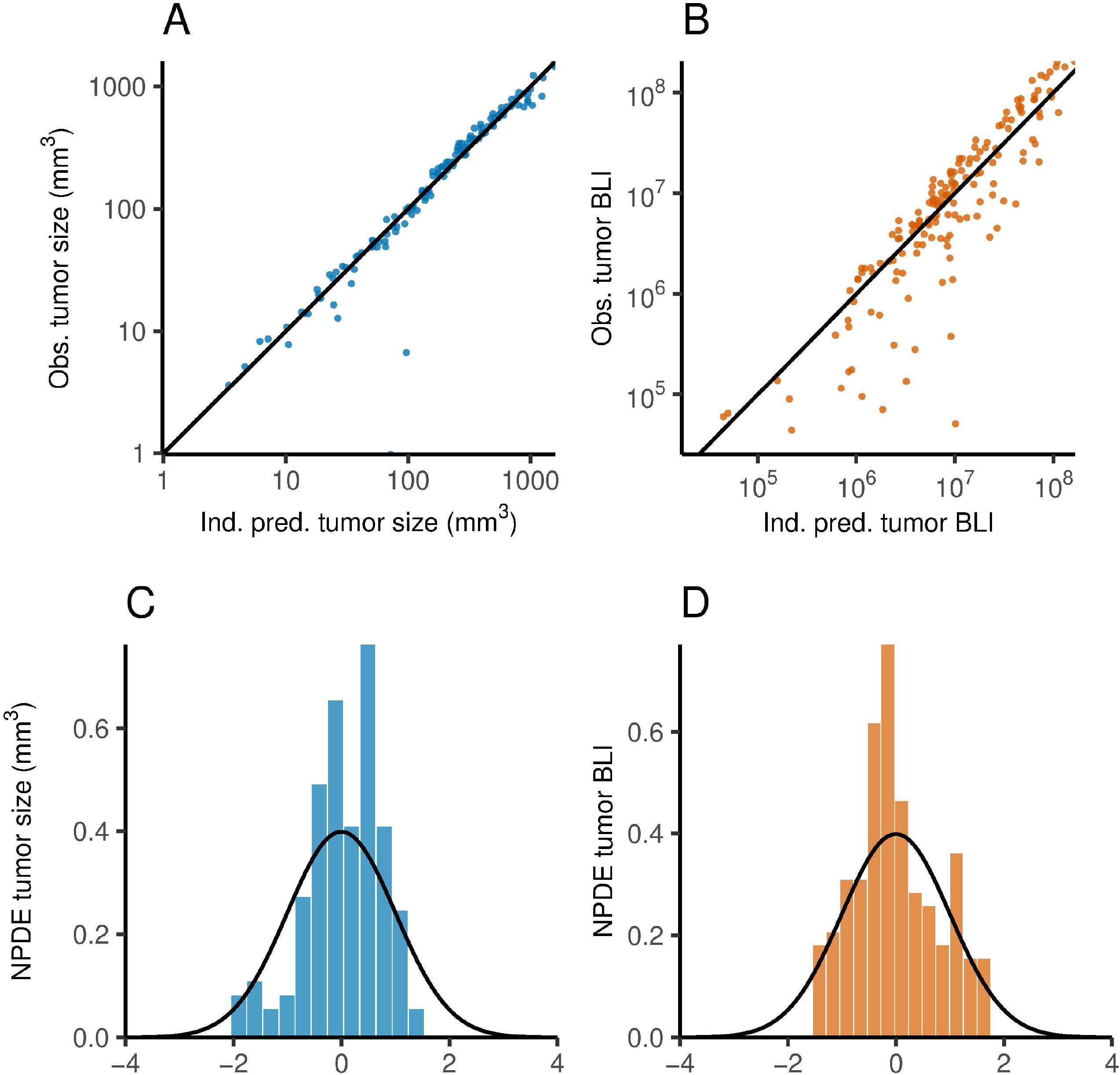
Diagnostic plots of the mathematical model. Observed vs. individual predicted tumor size and distribution of NPDE for tumor size (A and C) and tumor bioluminescencce (B and D), respectively. Observed tumor size and bioluminescence are correctly described by the model (A and B). The Normalized Prediction Distribution Errors (NPDE) do not display model misspecifications (C and D).

Population-level parameter estimates are well defined (Table 1). Individual parameter estimates exhibit substantial variability, with greater variability observed for BLI dynamics than for TS dynamics. The three ρ_50_ parameters, which quantify the TS/K ratio associated with half-maximal influence, are all greater than 1 at both the population and individual levels. Strong correlations between the ρ_50_ parameters were quantified. The γK and γL parameters take high estimated values, whereas γG is lower by comparison. No significant inter-individual variability was identified for these sigmoidicity parameters (Figure 5).

**Table 1:**
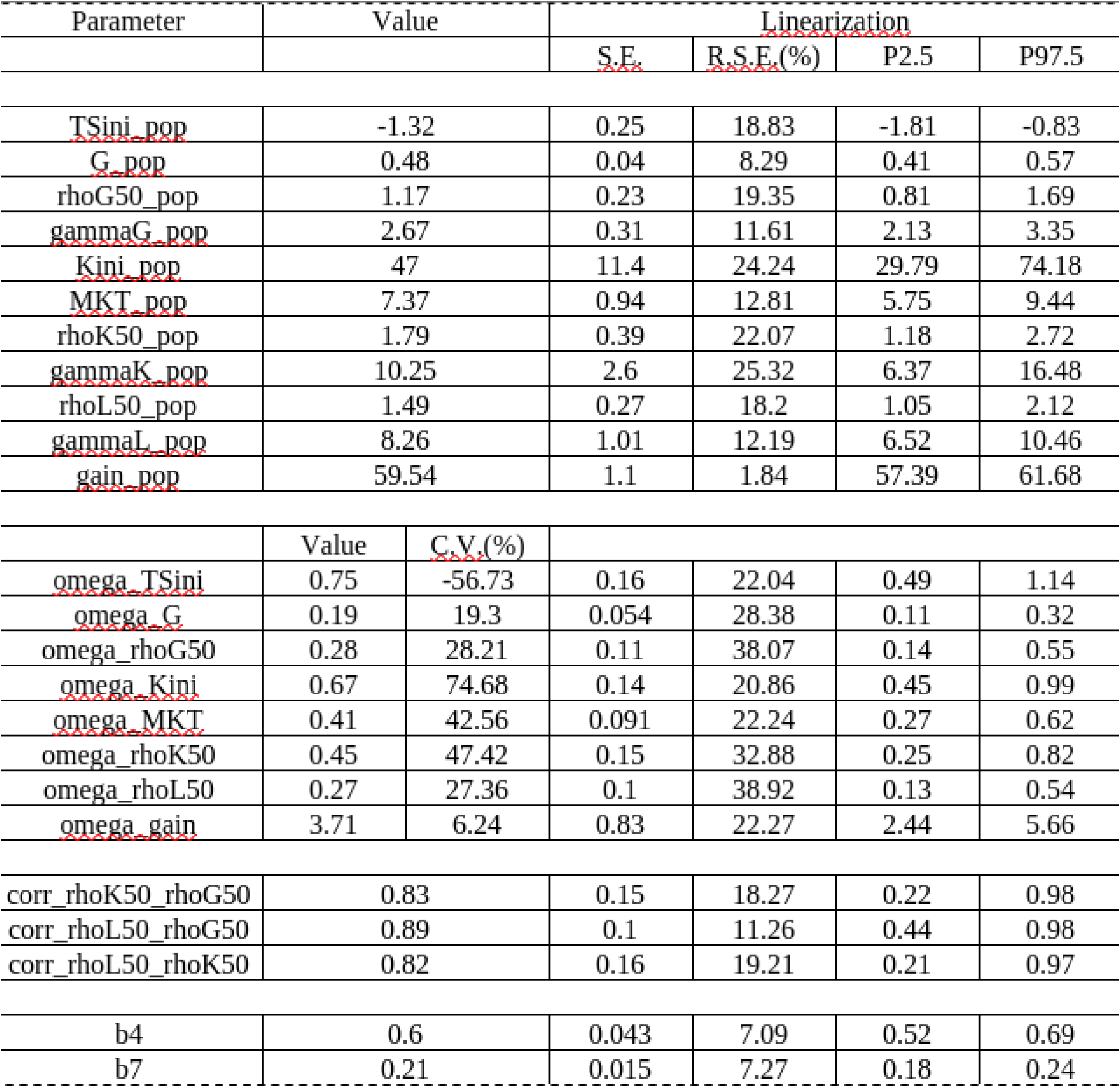
Estimated population parameter of the mathematical model.

**Figure 5.**
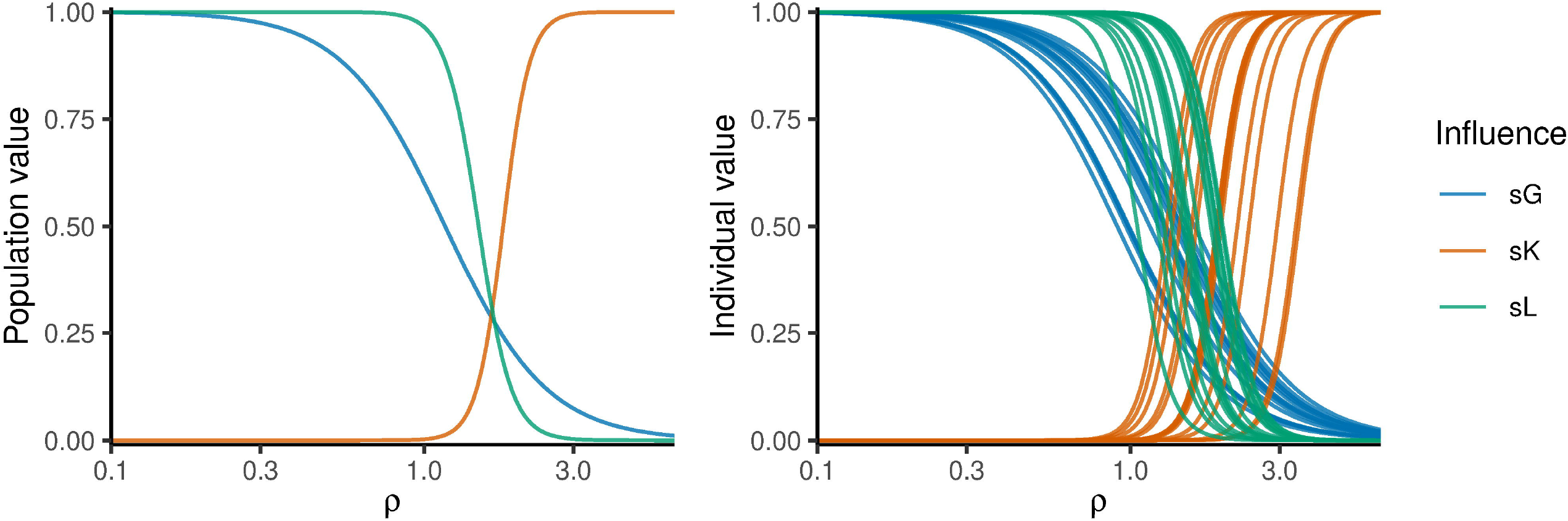
Simulations of the mathematical model. Population (left) and individual (right) simulations of the influence on growth rate sG (blue), on vascular capacity sK (red) and on bioluminesence signal sL (green) in function of the ratio ρ (TS/K). When ρ is low (vascular supply exceeds tumor size), tumor growth rate and bioluminescence are maximal, and vascular supply production is not stimulated. When ρ is high, tumor growth rate and bioluminescence are reduced, while vascular supply production is fully stimulated.

### Hypoxia evaluation

The percentage of pimonidazole appeared to trend with the estimated initial tumor size across the four subjects analyzed. The mouse with the lowest pimonidazole level corresponded to the largest estimated tumor size (10^TS_ini = 0.33 mm^3^), whereas the two mice with higher pimonidazole levels exhibited two of the three smallest tumor size values (<0.01 mm^3^). The mouse with a median pimonidazole percentage presented an intermediate tumor size (Figure 6).

**Figure 6.**
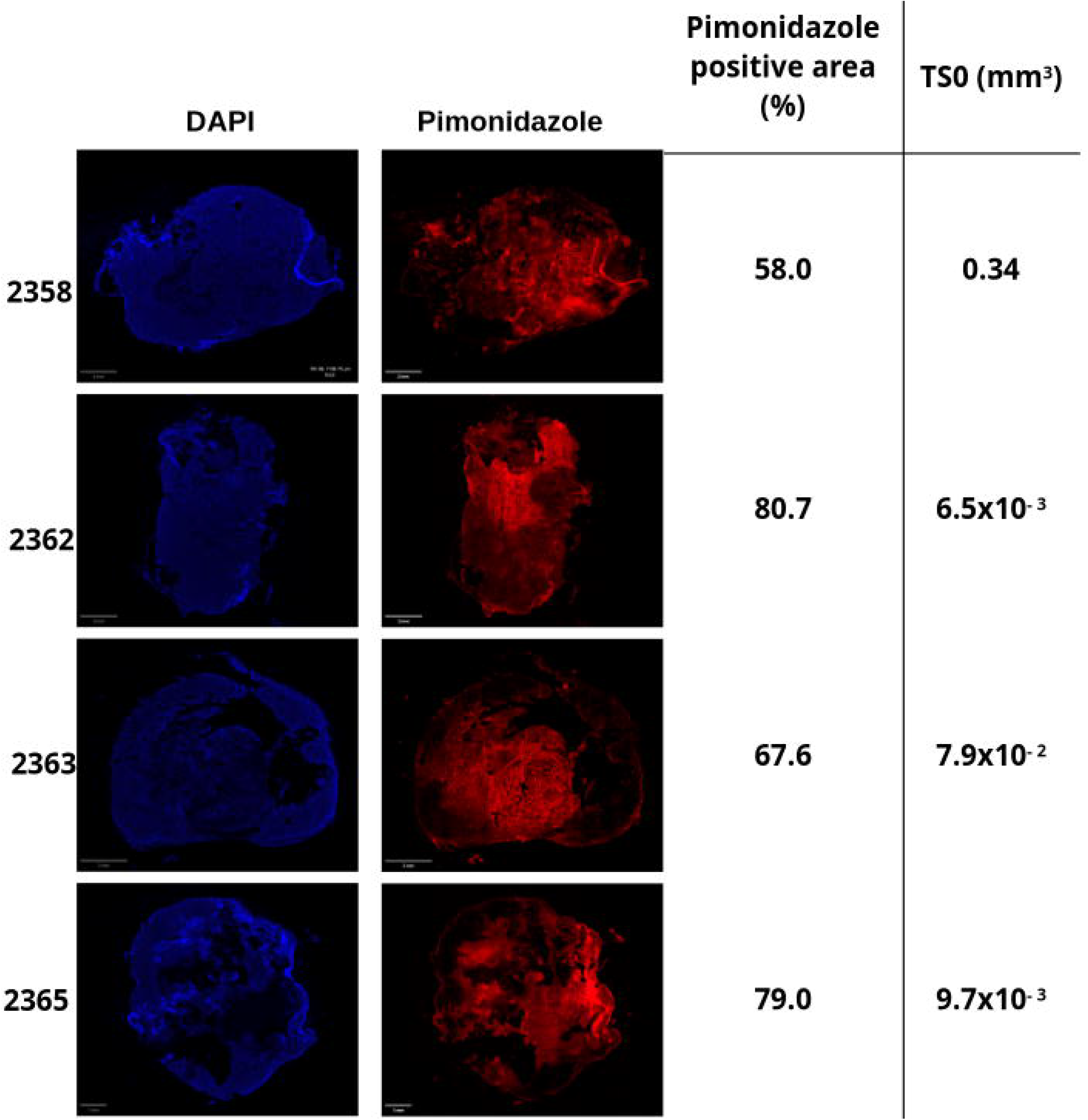
Tumor hypoxia and calculated initial tumor size. Representative images showing hypoxic regions in tumors from four mice, detected by pimonidazole (red) and DAPI (blue) staining. Scale bar, 1 mm. Quantification of the hypoxic tumor fraction was performed over the entire tumor area. TS0 denotes the model-predicted initial tumor size for each mouse.

## Discussion

In this study, we evaluated mammary tumor growth without treatment, in a syngeneic and orthotopic mouse cancer model using female BALB/cJ immunocompetent mice^14^. Three complementary non-invasive modalities were employed: caliper measurements for external tumor dimensions, ultrasound imaging for internal volumetric assessment, and bioluminescence imaging (BLI) for real-time monitoring of viable tumor cells. Tumor volume increased consistently according to both caliper and ultrasound imaging measurements, reflecting progressive tumor expansion. However, BLI signals did not correlate with volumetric measurements, initially decreasing and later increasing over time despite continued tumor growth. This discrepancy highlights a key limitation of BLI interpretation in the context of tumor hypoxia, a common feature of solid tumors that significantly impacts both biological behavior and imaging accuracy^15,16^.

Caliper measurements, while simple and non-invasive, provide a rapid estimate of tumor size through external palpation and calculation of ellipsoidal volume. However, they are prone to overestimation and variability due to irregular tumor shapes, inclusion of surrounding soft tissue or edema, and subjective measurement bias (Figure 1)^17^. Ultrasound imaging, in contrast, offers improved accuracy of volume estimation. It enables three-dimensional visualization and direct measurement of tumor boundaries, outperforming calipers in reproducibility, and correlation with actual tumor mass^18,19^.

The observed correlation between pimonidazole staining and estimated initial tumor size suggests that TSini - although a virtual parameter derived from the model - may reflect the level of tumor hypoxia. These findings indicate that, for the same number of injected cells, variations in the local hypoxic environment could lead to different estimates of the initial tumor size. Given the small sample size, this relationship should be interpreted cautiously, remains hypothetical, and requires further investigation.

However, such multi-modal imaging demands specialized equipment and expertise, limiting accessibility, and requires advanced quantitative analysis, and a high total cost. In our experiments, ultrasound imaging aligned with caliper measurements for overall volume trends (data not shown) but additionally revealed internal heterogeneity not captured by calipers. BLI, by contrast, provides a sensitive readout of viable tumor burden; however, the luciferase-luciferin reaction is oxygen-dependent, requiring molecular oxygen. In hypoxic tumor regions, photon output is reduced, potentially underestimating viable cell populations^12^. In our study, BLI kinetics frequently decreased following initial growth phases, while caliper and ultrasound imaging measurements confirmed ongoing expansion. Previous studies indicate that severe hypoxia can reduce bioluminescent signals by up to 50%, which may confound interpretation of tumor progression and treatment efficacy^11^. This limitation was found in our data, where BLI kinetics often showed a decrease after initial growth phases, while caliper and ultrasound imaging confirmed ongoing expansion.

To quantitatively describe the relationship between tumor growth, hypoxia, and BLI, we developed a mathematical compartmental model incorporating tumor volume, cell proliferation, and vascularization parameters. The model provides a robust framework for TS and BLI dynamics. Individual parameter estimates exhibited substantial variability, particularly for BLI, reflecting heterogeneity in tumor physiology, microenvironment, and vascularization. The ρ_50_ parameters were consistently greater than 1, with strong correlations among them, indicating that relatively high TS/K ratios are required to reach half-maximal influence. The γK and γL parameters exhibited high values, whereas γG was comparatively lower, suggesting a pronounced influence of vascularity on tumor growth and BLI. No significant inter-individual variability was observed for these sigmoidicity parameters, indicating conserved vascularity-dependent dynamics across the population.

Most existing tumor growth models are tumor growth inhibition models (TGI) models focusing on drug effect quantification. Few of these models account for hypoxia^20^ but don’t distinguish drug effect from vascular supply limitations. The model proposed by Ribba *et al*. accurately describes tumor growth in the absence of treatment; however, it does not incorporate hypoxic oscillations^21^. In contrast, the integrative model presented here incorporates latent vascular capacity modulation as a central structural component, governing tumor cell maturation, growth, and BLI properties. This feature allows a more mechanistic representation of tumor dynamics beyond standard TGI approaches.

Nevertheless, the model has limitations. It does not include tumor elimination or mechanical constraints on growth, resulting in theoretically unbounded size increases predictions. While this did not impact the current results due to the limited observation period, it would need to be addressed in longer-term studies. The model also does not account for variations in luciferase stability or signal attenuation, which could influence BLI measurements. The incorporation of these factors in future modeling studies could enhance the accuracy of TS growth predictions. The perspectives could be - the characterization of how genetic polymorphisms of vascular pathway–related actors influence tumor growth dynamics. - Or the optimization of measurement timing to capture key tumor growth modulations and vascularization events more accurately. Overall, this integrative model provides a quantitative framework for describing tumor growth and BLI dynamics, highlights the role of vascularity, and identifies sources of inter-individual variability. Future refinements incorporating additional biological and technical factors could further enhance its utility for interpreting tumor progression and evaluating preclinical interventions.

## Author contributions

SR, AL and SC designed the experiments; JS, SN and SL performed the experiments and contributed to the data collection; NA, DT, SR and SC contributed to the interpretation of data; NA and SC performed all data integration and modelling analysis; NA, SR and SC wrote the original drafts; NA, DT, JS, SN, SL, AL, SR and SC reviewed and edited the manuscript; SR contributed to the funding acquisition and project conceptualization; SC performed project supervision.

## Acknowledgement

The authors acknowledge H2P2 platform for technical assistance for immunohistochemistry analysis.

## Data availability

The datasets used and/or analyzed during the current study are available from the corresponding author on reasonable request.

